# The respiratory phase modulates task-related neural representations of visual stimuli

**DOI:** 10.1101/2024.12.03.626336

**Authors:** Lisa Stetza, Lena Hehemann, Christoph Kayser

## Abstract

We investigate how respiration influences cognition by examining the interaction between respiratory phase and task-related brain activity during two visual categorization tasks. While prior research shows that cognitive performance varies along the respiratory cycle, the underlying neurophysiological mechanisms remain poorly understood. Though some studies have shown that large-scale neural activity reflecting changes in the excitation-inhibition balance is co-modulated with the respiratory cycle, it remains unclear whether respiration directly shapes the quality by which task-relevant sensory information is encoded. We address this gap by applying single-trial multivariate analyses to EEG data obtained in humans, allowing us to track how respiration modulates the sensory evidence in this neurophysiological signal. Confirming previous studies, our data show that participant’s performance varies with the respiratory phase prior and during a trial. Importantly, they also suggest that respiration directly influences the sensory evidence carried by parieto-occipital processes emerging around 300 to 200 ms prior to participant’s responses. Hence, respiration and sensory-cognitive processes are not only highly intertwined but respiration directly facilitates the representation of behaviourally-relevant signals in the brain.

## Introduction

The potential benefits of structured respiration for brain and bodily functions have been considered for centuries. Still, the precise interplay between respiration and cognitive functions as well as the underlying cerebral processes remain a matter of current research (Varga and Heck, 2017; Brændholt et al., 2023; Kluger et al., 2024; Madsen and Parra, 2024). While previous work shows that behavioural performance varies along the respiratory cycle, few studies have directly investigated the neurophysiological basis for a co-modulation of behaviour along the respiratory cycle. We here bridge this gap using single-trial multivariate analysis applied to EEG data derived during two visual tasks and show that respiration directly modulates the encoding of task-relevant visual representations in the brain.

Previous work has shown that participants tend to align their respiration to the series of experimental trials (Johannknecht and Kayser, 2022; Goheen et al., 2024). At the same time, trial to trial variations in respiratory phase are predictive of how well or how fast participants respond in perceptual or cognitive tasks (Flexman et al., 1974; Gallego et al., 1991; Huijbers et al., 2014; Arshamian et al., 2018; Perl et al., 2019; Zelano et al., 2016; Nakamura et al., 2018; Park et al., 2020; Johannknecht & Kayser, 2022; Mizuhara & Nittono, 2023; Braendholt et al., 2024). Furthermore, changes in motor performance and brain-muscle coupling along the respiratory cycle have been reported (Li and Laskin, 2006; Li and Rymer, 2011; Kluger and Gross, 2020; Park et al., 2020; Engelen et al., 2024). Hence, the converging evidence for a modulation of participant’s behaviour along the respiratory cycle is strong, but the underlying neurophysiological basis remains unclear.

The brain structures controlling respiration (e.g. the pre-Boetzinger complex) and those sensing the resulting changes in airflow or chest pressure are intricately connected with the limbic and autonomous nervous system (Del Negro et al., 2018; Ashhad et al., 2022; Krohn et al., 2023). As a result, direct neural feedback about the current respiratory state is potentially widely available in the brain. Indeed, intracranial recordings have shown that the respiratory rhythm entrains faster oscillations beyond the olfactory cortex, presumably by coordinating long-range communication within and across brain regions (Zelano et al., 2016; Tort et al., 2018; Herrero et al., 2018). Such a brain-wide coordination of activity along the respiratory cycle is also evident in neuroimaging studies. Using MEG, Kluger and colleagues showed that the prominence of both rhythmic and aperiodic activity varies along the respiratory cycle in different parts of the brain (Kluger & Gross, 2021; Kluger et al., 2023). Such large-scale changes in brain activity may possibly reflect changes in the excitation-inhibition balance coordinated with respiration (Kluger et al., 2021).

While those studies show that information about the respiratory state is potentially widespread in cortical and subcortical regions, it remains unclear which of those neurophysiological signatures reflect or directly mediate the changes in behavioural performance along the respiratory cycle. For example, a co-modulation of behavioural performance with the respiratory cycle could originate from changes in brain-muscle coordination, from changes in attention, or from very specific changes in the task-relevant sensory representations. While changes in brain-muscle coordination and attention-related processes along the respiratory cycle have been shown (Kluger & Gross, 2020; Kluger et al., 2021; Engelen et al., 2024) it remains unclear whether respiration also modulates specific cerebral representations of the sensory information used in a task. While a study on eye blink conditioning suggests that respiration can indeed modulate sensory evoked potentials (Waselius et al., 2022), it remains unclear whether this translates to purely perceptual tasks and whether respiration modulates the strength by which cerebral processes reflect external sensory information. Based on human EEG data obtained in two visual categorization tasks we here probe for such specific changes in cerebral representations along the respiratory cycle. Based on multivariate classification we extracted the neurophysiological signatures of task-relevant information and probed which of those are modulated by respiration in a manner that is also predictive of behaviour.

## Methods

### Participants

In this study, 27 adult volunteers participated after providing informed consent. All had normal hearing and vision and were compensated for their time. An age limit was set to 40, but detailed demographic data was not collected; overall, the group of participants consisted of young university students. Participants were informed about the procedures and devices used in the experiment but were not explicitly told that the main focus was to study the influence of respiration on task performance and brain activity, similar as in previous work (Johannknecht & Kayser, 2022). Participants were asked to breathe normally through their nose. We cannot rule out that some participants switched between oral and nasal respiration while performing the task. All procedures were approved by the ethics committee of Bielefeld University.

### Experimental setting and data collection

The experiments were performed in an electrically shielded and sound-attenuated booth (Desone, Germany). The stimuli were projected (LG HF65, LG electronics) onto a 2x1m screen (Screen International Modigliani) about 1m away from the participant. Stimulus presentation was controlled via Psychophysics Toolbox (Version 3.0.14) using MATLAB (Version R2017a, The MathWorks, Inc., Natick, MA) and synchronised with TTL pluses to an EEG recording system (ActiView, BioSemi). Participants responded using a computer keyboard.

Respiration data was recorded using a temperature-sensitive resistor (Littelfuse Thermistor No. GT102B1K, Mouser electronics) that was attached to a modified single-use clinical oxygen mask (Johannknecht & Kayser, 2022). The resistor detects the temperature changes in the respiratory airflow associated with inspiration and expiration. The corresponding voltage changes across the resistor were amplified and recorded using the analogue port of an ActiveTwo EEG system (BioSemi BV; Netherlands). Electrophysiological data were recorded using a 128 channel EEG system at a sampling rate of 1024 Hz, using Ag-AgCl electrodes mounted on an elastic cap (BioSemi BV; Netherlands). Electrode offsets were kept within ±25 mV. The EOG data were recorded using four electrodes placed at the side and underneath each eye.

### Experimental tasks

Participants performed two tasks involving the discrimination of emotions and the discrimination of visual shapes. Both tasks manipulated the task difficulty in a parametric manner. They were each presented in separate blocks, with the first three blocks of each session featuring visual shape discrimination and the last three blocks emotion discrimination.

### Emotion discrimination

This task was similar to one used in a previous study on respiration (Johannknecht & Kayser, 2022). Participants were presented with images showing happy or angry facial expressions and had to categorise these emotions. The stimuli were derived from the Dynamic FACES database (Holland et al., 2019), from which we selected frames in the transition from an emotional expression to a neutral expression to provide two levels of emotional expression (strong and intermediate). In total, we selected images from 24 individuals balanced across age and sex (3 age cohorts, 2 sexes). For each of those individuals we selected 5 images: a strong and intermediate expression of each emotion and a neutral expression. In the experiment, these images were presented at an angle of 15° x 12° (vertical x horizontal) for 16 ms and participants were instructed to indicate as fast as possible which emotion they perceived. Participants completed 3 blocks of 160 trials (1200-1500 ms inter-trial interval; 400-1000 ms fixation interval) with a balanced presentation of individuals, emotions and levels within and across blocks.

### Visual shape discrimination

Participants were presented with (semi-)ambiguous images of a Necker-like cube and had to discriminate between the two possible orientations of the cube (i.e. to indicate which side was in front). The images subtended 11° x 9°, were constructed from light grey lines on a dark grey background and were presented for 50 ms. To manipulate task difficulty, we systematically varied the luminance of the lines that formed the front side of the cube in 5 levels, aiming for a gradual transition from the perception of the left-hand side to be in front to the right-hand side to be in front (Fig. 1). The intermediate level presented a fully ambiguous cube. Participants were instructed to answer as fast as possible with previously assigned keys. As in the emotion paradigm, participants completed 3 blocks of 160 trials (800-1200 ms inter-trial interval, 1100-1500 ms fixation interval) with a balanced presentation of cube orientation and ambiguity within and across blocks.

**Figure 1:**
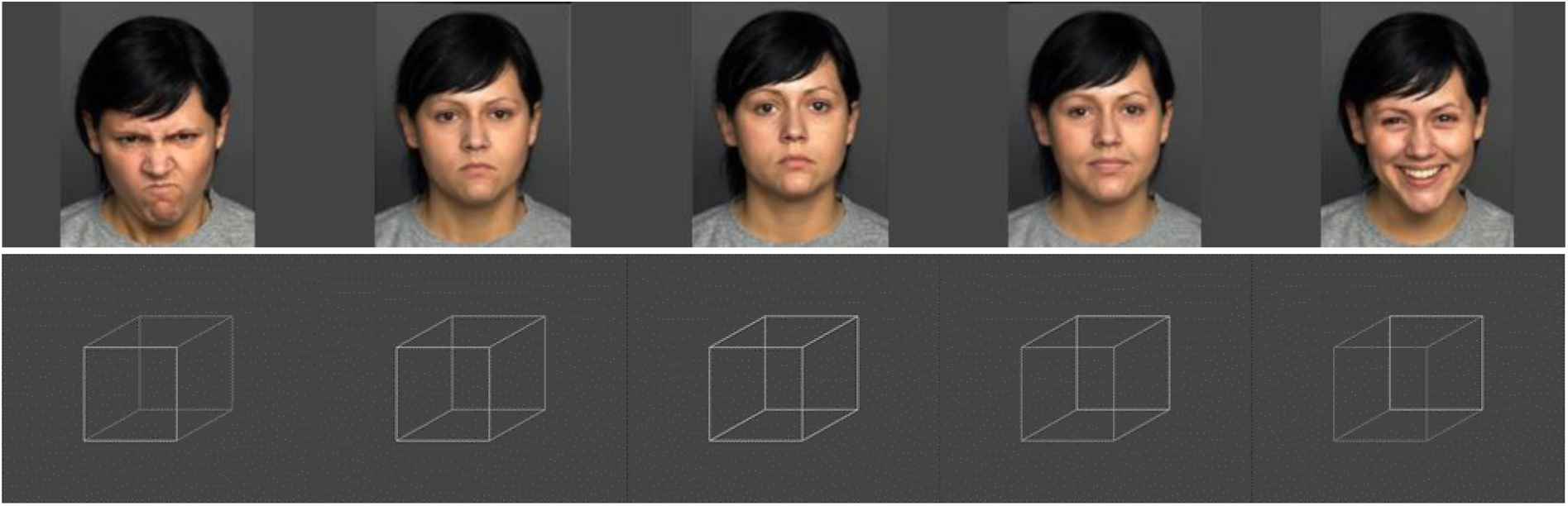
Example stimuli. (A) Exemplary images showing emotional expressions transitioning from angry to happy used in the emotion discrimination task (derived from and shown with permission of the Dynamic FACES database; Holland et al. 2019). Note that these particular images were not used in the experimental trials and are shown here as they are part of the images allowed for displayment. (B) Images of the (semi-) ambiguous cubes ranging from a clearly left-oriented variant to a clearly right-orientated variant used in the visual shape task.

### Analysis of respiratory data

The raw signals were filtered (3rd order Butterworth filters; high pass 0.03 Hz, low pass 6 Hz) and resampled at 100 Hz using the FieldTrip toolbox (Oostenveld et al., 2011) in Matlab (Mathworks Inc., Natrick, MA; version R2022b). The Hilbert transform was used to identify individual respiratory cycles by determining local peaks based on the respective phase variable. Individual cycles were defined based on the data in windows of 7 seconds around each peak, whereby peaks were included if the z-scored value exceeded a level of z=0.5. The inspiration period was defined as a continuous period with a positive slope preceding the local peak whereas expiration was defined as the negative slope following the peak. Interruptions in these periods shorter than 500 ms were interpolated. As a result, each respiratory cycle was split into inspiration and expiration phases, though for some cycles short exhale pauses were classified as third state and not analysed (Noto et al., 2018). We compared the distribution of participant-wise respiratory cycles based on their mean square distance along time and removed cycles whose mean squared distance was greater than 3 standard deviations from the centroid as atypical cycles from subsequent analysis.

To link respiratory signals to behaviour we defined the phase of each respiratory cycle as a linearly increasing variable from the beginning to the end of inspiration (defined as angle from 0 to pi) and subsequently as linearly increasing from the beginning to the end of expiration (defined as pi to 2*pi). This phase variable scales linearly in time within each inspiration or expiration period, and has two clearly interpretable time points (0/2pi, and pi). To test for an alignment of respiration to the experimental trials we calculated the phase coherence for each participant as follows (Johannknecht & Kayser, 2022; Park et al., 2020): we converted the respiratory phase to a complex valued number, averaged these across trials and took the resulting vector length. We established a baseline estimate of the phase coherence expected under the null-hypothesis of no alignment of respiration to the experimental paradigm by computing for each participant a distribution of phase coherences after randomising (2000 times) the alignment of respiration to the experimental trials by randomly shifting these.

### Modelling behaviour based on respiratory phase

To probe for a statistical relation between respiration, task-related variables and behaviour we used linear mixed effect models. As predictors we included a variable reflecting the parametric manipulation of task difficulty (stimulus level), the category of the stimulus (e.g. emotion), and modelled respiration using the sine and cosine transformed trial-wise respiratory phase derived at particular time points of interest. These time points were derived in an epoch around either the time of stimulus onset or the time of each trial’s response (sampled in 300ms steps; Fig. 2C&D). We also included a participant-wise random effect of trial number to capture potential effects of fatigue or training-on-the-task and a random offset for each participant. For reaction times (RT) the model was as follows, with RespSine and RespCosine denoting the transformed phase of the respiration defined at a particular time point of interest:

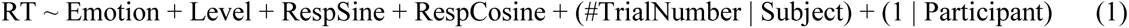

**Figure 2:**
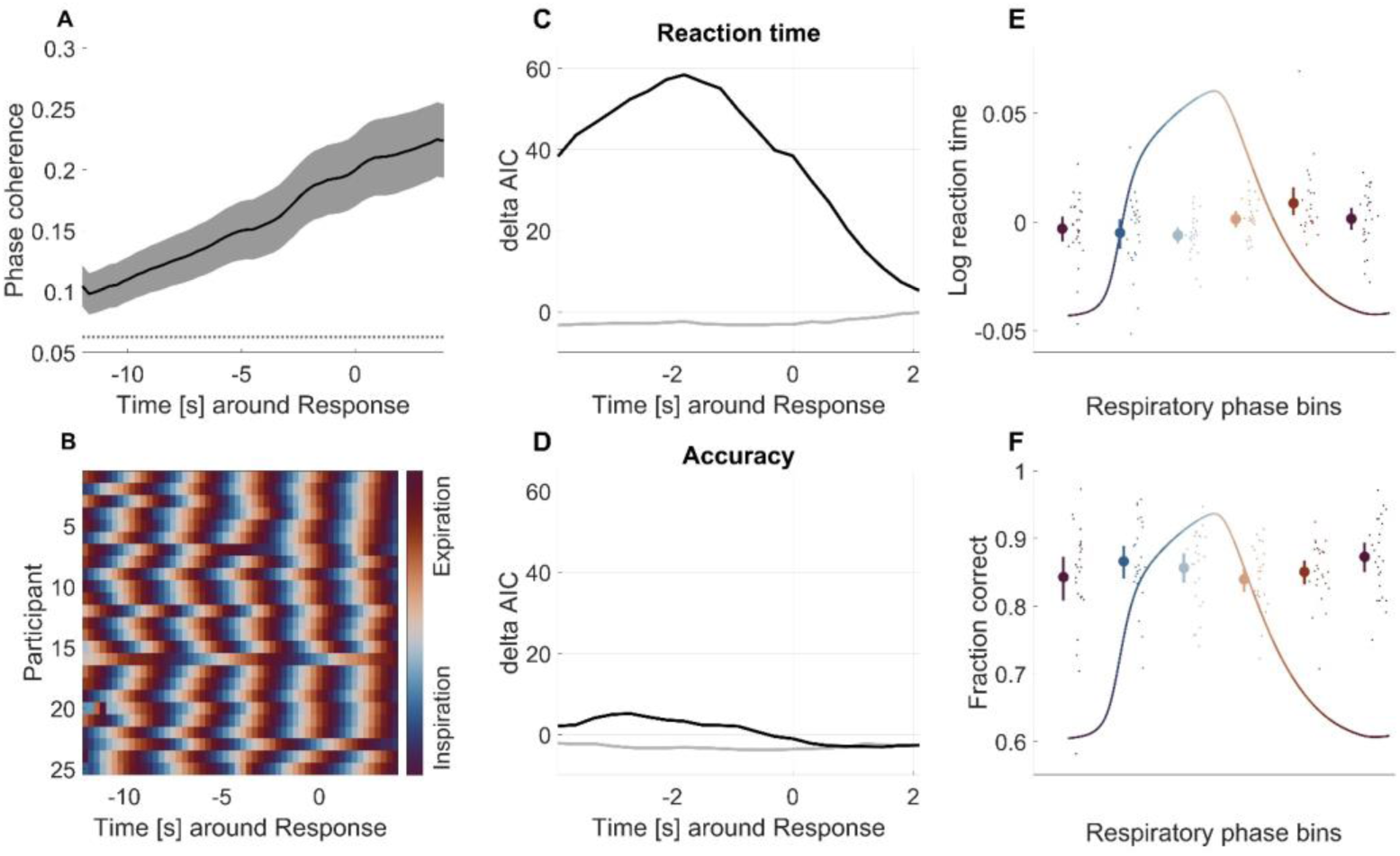
Behavioural and respiratory data for emotion discrimination. (A) Consistency of the respiratory phase across trials relative to the response times (mean, s.e.m. across participants); dotted line indicates significance threshold (p<0.01). (B) Trial-averaged respiratory phase for each participant. (C) AIC differences for a predictive effect of respiratory phase on reaction times; the grey line shows the effect for respiration aligned to stimulus onset, the black line for respiration relative to response times. Positive values indicate an effect of the respiratory phase. (D) Same for accuracy. (E) Average normalised reaction time across participants for trials with a given respiration phase (divided into six bins; at the time of response; mean, ± s.e.m.). Dots indicate individual participants. (F) Same for accuracy.

The reaction times were square-root transformed before entering the model. The models for reaction times assumed Gaussian variables and used a linear activation function, the corresponding model for accuracy used a logistic activation function and assumed binomial variables. For this analysis, we excluded trials with overly short reaction times (<200 ms) or excessively long reaction times (based on the median absolute deviation criterion) and those (rare) trials for which the respiratory phase could not be defined.

We derived two statistical measures of the relevance of each factor of interest in these models: first, we determined the single trial predictive power of each factor by comparing the full model (eq. 1) with a model not including the respective factor. This comparison was implemented by computing the difference in the Akaike information criteria (AIC) of the two models (Wagenmakers & Farrell, 2004). Based on the notion of relative model probabilities (Akaike weights) we considered an AIC difference > 13.8 as evidence for a meaningful effect; this corresponds to a probability of 99% or higher of the model including the factor of interest to have more explanatory power.

Second, we tested for a relation between the dependent variable and each factor by assessing the statistical significance of the respective predictors themselves. For emotion and level these were based on the respective t-statistics, for respiration this was implemented by comparing the vector strength of the combined sine and cosine respiratory predictors to surrogate data (Kluger & Gross, 2021; Johannknecht & Kayser, 2022). We shuffled the trial-wise relation of the predictors and the dependent variables, computed the corresponding model and extracted the respective vector strength, repeating these steps 6000 times. Based on the surrogate distribution we derived the probability of the actual values to exceed chance. To visualise the relation between respiratory phase and reaction times (response accuracy; e.g. Fig 2E&F), we grouped trials by binning the respiratory phase into six equally spaced bins and derived the average reaction time (% correct responses) for each bin and participant.

### EEG pre-processing

The data were bandpass filtered between 0.6 Hz and 90 Hz, resampled to 200 Hz and epoched from 0.8 s to 1.8 s around stimulus onset. Bad channels were interpolated where necessary (mean ± SD of interpolated channels per participant; shape: 0.7±0.2, emotion: 0.8±0.2). Potential artefacts were removed using independent component analysis (ICA), computed across all blocks. We removed ICA components that reflect eye movement artefacts, localised muscle activity or poor electrode contacts (mean ± SD removed components per participant; shape: 9.8±0.9, emotion: 9.6±1). These were identified as in our previous studies (Kayser et al., 2017; Grabot and Kayser, 2020) following definitions provided in the literature (O’Beirne and Patuzzi, 1999; Hipp and Siegel, 2013). Trials with amplitude exceeding 250mV were rejected. For the subsequent analysis of EEG data, we also excluded trials with excessive reaction times or an undefined state of respiration (see above). In addition, we excluded those trials with the arbitrary stimulus category, as for those the category of the stimulus and the correct response are not defined. This left on average 354±4.0 (mean±SEM) trials for emotion and 363 ± 2.2 (mean±SEM) trials for the shape paradigm per participant.

### Linear discriminant analysis

The main analysis probed whether respiration modulates the representation of task-related sensory representations in the EEG. To extract these, we used single-trial multivariate linear discriminant analysis (LDA) to identify neural signatures of the emotion category or the visual shape (Park & Kayser, 2021; Guggenmos et al., 2018; Parra et al. 2005). We then probed whether the trial-wise neural evidence about the stimulus category carried by these signatures was modulated by respiration and whether this evidence is also related to participants’ responses. For this the EEG data were filtered between 0.8 and 30 Hz (3^rd^ order Butterworth filter). The LDA was applied to the data aligned to stimulus onset (or response times) in sliding windows of 60 ms duration, moving in 5ms time steps. The regularisation parameter was set to 0.1 as in previous work (Park & Kayser, 2021). The LDA classification performance was determined using the area under the curve (AUC) of the receiver operating characteristic obtained from six-fold cross-validation. Forward models for each LDA component were derived as the normalised correlation between the discriminant component and the EEG activity (Parra et al., 2005).

Based on the group-level forward models we established LDA components of interest within the time window of significant classification performance (see below). For this, we relied on the clustering of forward models in time to group neighbouring components reflecting similar neurophysiological origins. For each cluster, we determined the peak time point of the group-level classification performance and used these time points to define components of interest; for the emotion paradigm these peak time points were 237 ms and 452 ms after stimulus onset and 360 ms and 245 ms prior to the response; for visual shape 267 ms and 382 ms after stimulus and 440 ms and 255 ms preceding the response. For subsequent analysis, we derived the trial-wise projection of each LDA component of interest. Practically, for each component and participant, we first defined the peak AUC for this participant in a time window of ±100 ms around the respective component time indicated above. This was done to define a participant-specific LDA component that has maximal sensitivity to the stimulus in the relevant time period. We then used the LDA weights associated with this participant-specific component and derived the trial-wise projection of the component activity during the entire trial (Fig. 4, upper panels). The distance of this projection from the LDA decision boundary reflects a measure of the ‘stimulus evidence’ available at each moment in time (Franzen et al., 2020; Philiastides et al., 2014).

We then asked whether this stimulus evidence is modulated by respiration. For this we grouped trials according to respiratory phase, using 6 equally-spaced phase bins, and averaged the stimulus evidence within each phase bin. We then used linear modelling to probe whether and when stimulus evidence is modulated by respiratory phase. We implemented this analysis for each of the four components of interest. For the two components defined based on stimulus-aligned classification, we defined the respiratory phase at different time points relative to stimulus onset; for the two components defined based on the response-aligned classification we defined the respiratory phase relative to the trial-wise response time. Note that this analysis effectively comprises two time dimensions: the time within each trial relative to the moment at which the LDA component was defined, and the time (relative to stimulus or the response) at which the respiratory phase was defined (c.f. Fig. 4).

### Statistical analysis of EEG data

Group-level inference on the significance of the classifier performance was performed using a randomization procedure and cluster-based statistical enhancement controlling for multiple comparisons along time (Nichols and Holmes, 2002; Maris and Oostenveld, 2007). To test the group-average AUC we shuffled the sign of the true single-participant effects (the signs of the chance-level corrected AUC values) and obtained the respective distribution of the group average over 4000 randomizations. The first-level effects were thresholded based on the 99.9th percentile (i.e., p=0.001) of the randomization distribution and significant time bins were clustered based on a minimal cluster size of six and using the maximum as cluster statistics. We report significant clusters at a second-level significance of p<0.01.

To test for an effect or respiratory phase on stimulus evidence, we modelled the phase-binned stimulus evidence against the sine- and cosine predictors of respiratory phase (and an offset), separately for each time point of respiration and the time within each trial (relative to the respective peak time of the LDA component). As for the behavioural data, we derived two measures of statistical significance: first, we compared the predictive power of a model including respiration to one without and derived the associated AIC difference similar as for the behavioural data. This is shown in Fig 4, lower panels along both time axes. We considered AIC differences as meaningful that correspond to a probability of the model including respiration to explain the data better at p<0.0001. We used such a stringent criterion given the multiple comparisons performed across time bins. Second, we relied on permutation statistics to probe for the significance of the vector sum of the respiratory predictors. Given that the respiratory phase is redundant over the time course of respiration, we averaged the vector length of true and surrogate effects over this time axis to reduce the number of time points in the analysis. We then used a clustering procedure to probe for significant effects along the time axis of the LDA component, correcting for multiple comparisons along this dimension (using a first-level threshold of p<0.001; a minimal cluster size of 6 and the max-sum as clustering statistics, using 6000 permutations). Overall both approaches yielded converging results in that the same components around the similar classifier time points reached the respective thresholds of statistical significance.

The above analysis revealed specific time windows for some of the LDA components at which the stimulus evidence carried by these is significantly modulated. We asked whether this stimulus evidence is also related to behaviour, by comparing linear regression models, one incorporating the stimulus evidence of each identified LDA component to predict behavioural performance, the other neglecting stimulus evidence.

## Results

### Emotion categorization varies with respiratory phase

We first focus on the emotion paradigm and report results for the shape task below. During the categorization of emotions, the reaction times varied with the emotion category (p<10^-5^, t=9.3, deltaAIC=84) and the level by which the emotion was expressed (p<10^-5^, t=16.0, deltaAIC=251). Participants reacted faster when the emotional expression was stronger and were faster for happy faces (see Table 1). The response accuracy also varied with the emotion (p=0.0013, t=3.22, deltaAIC=8) and the level of expression (p<10^-5^, t=16.0, deltaAIC=277; Table 1).

**Table 1:** Averaged reaction times in milliseconds (mean and standard error) across participants for each stimulus category and level of difficulty.

| Strong anger | Intermediate anger | Intermediate happiness | Strong happiness |
| --- | --- | --- | --- |
| $809 \pm 4$ | $834 \pm 4$ | $833 \pm 4$ | $752 \pm 3$ |

| Left disambiguated | left intermediate | right intermediate | right disambiguated |
| --- | --- | --- | --- |
| $692 \pm 3$ | $707 \pm 3$ | $689 \pm 3$ | $654 \pm 2$ |

In line with previous work, participant’s respiratory phase was systematically related to the experimental trial, both within-trials for each participant and across participants. Figure 2A displays the between-trial consistency of respiratory phase, which was above chance level (at p<0.01). Hence for each participant the respiratory phase featured a consistently similar phase across trials. Figure 2B displays the trial-averaged respiratory phase angle for each participant. This shows that most participants exhibit a similar average phase around the time of each trial, showing that respiration is structured both within and across participants.

Despite this alignment of respiration to the experimental paradigm, there remains considerable trial-to-trial variability in respiratory phase and this variability is significantly predictive of trial-to-trial fluctuations in behaviour. We established this co-modulation of behaviour with respiration by comparing the predictive power of statistical models including or excluding respiration as predictor. Figure 2C shows the respective AIC differences for a relation of respiratory phase and reaction times tested using the respiratory phase at different time points prior to stimulus onset (grey curve, sampled in 300ms steps) or prior to the trial-wise response (black curve). For reaction times this co-modulation with respiration was very strong (peak deltaAIC= 58.4) and peaked for the respiratory phase about 2 seconds prior to the response. Relative to stimulus onset, this modulation was not present (peak deltaAIC = -0.2). For response accuracy (Fig. 2D), we did not find evidence for a co-modulation with respiration (response-aligned: peak deltaAIC=5.2; stimulus-aligned peak deltaAIC = -2.2). Also, we found no evidence for interaction between respiration and emotion intensity in predicting reaction times (deltaAIC=-3.0) or accuracy (deltaAIC = -6.5).

### EEG components sensitive to emotions

We used a linear discriminant analysis (LDA) to identify EEG components sensitive to the emotional expressions. Figure 3 shows the classifier performance, for the EEG data aligned to stimulus onset and when aligned to the trial-wise response times. Classification performance was significant between 187 ms and 697 ms after stimulus onset and -485 to 125 ms prior to responses (cluster-based permutation tests; p<0.001). Based on the group-level forward models of the participant-wise classifiers we identified two main EEG components of interest, separately for each alignment of the data. The corresponding forward models are shown in Figure 3 insets, the respective peak-times are 237 ms (termed ‘early’ in the following) and 452 ms (‘late’) for the stimulus-aligned data and -360 ms (again ‘early’) and -245 ms (‘late’) for the response-aligned data. The topographies of these components reflect the largely symmetrical relative engagement of fronto-occipital electrodes. Given that the typical reaction time in this paradigm was about 800ms (Table 1), the time of the late component in the stimulus-aligned data (452 ms) corresponds to about the same time in the trial as the time point of the early component in the response aligned data (-360ms). Hence, these two components may possibly reflect related neurophysiological processes.

**Figure 3:**
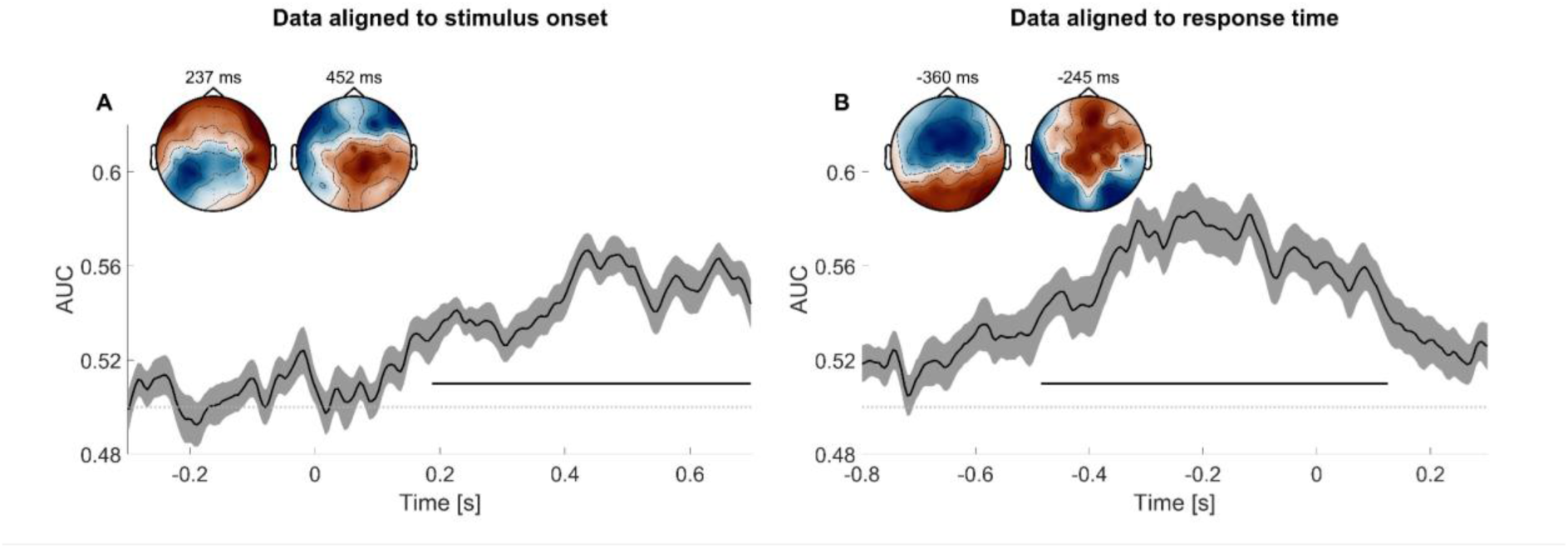
EEG components carrying evidence about the emotion category. (A) Classifier performance (AUC) for the data aligned to stimulus onset. Black line shows group average with the standard error as grey shade. Based on the LDA forward models we identified two components, the first peaking around 237 ms, the second around 452 ms after the stimulus onset. (B) Classifier performance for the data aligned to the response. Again two components were identified, one peaking 360 ms before the response, the other 245 ms before the response. The black horizontal lines indicate significant classifier performance (at p<0.001; cluster-based permutation test).

### Respiration shapes the behaviourally-relevant encoding of emotions

For each of the four components we derived the single trial time course of the respective stimulus evidence. Evidence here refers to the strength of the classifier prediction about the emotion presented on each trial (defined as the distance of the trial-wise classifier output and the respective decision boundary; Figure 4, upper panels). We then tested whether this stimulus evidence is significantly modulated by the respiratory phase. As for behaviour, we relied on the comparison of linear models, computing separate models for the respiratory phase defined at different time points in the trial and the EEG signal at different time points relative to the respective LDA component time. This revealed a significant and widespread modulation of emotion evidence by respiration for two components.

**Figure 4:**
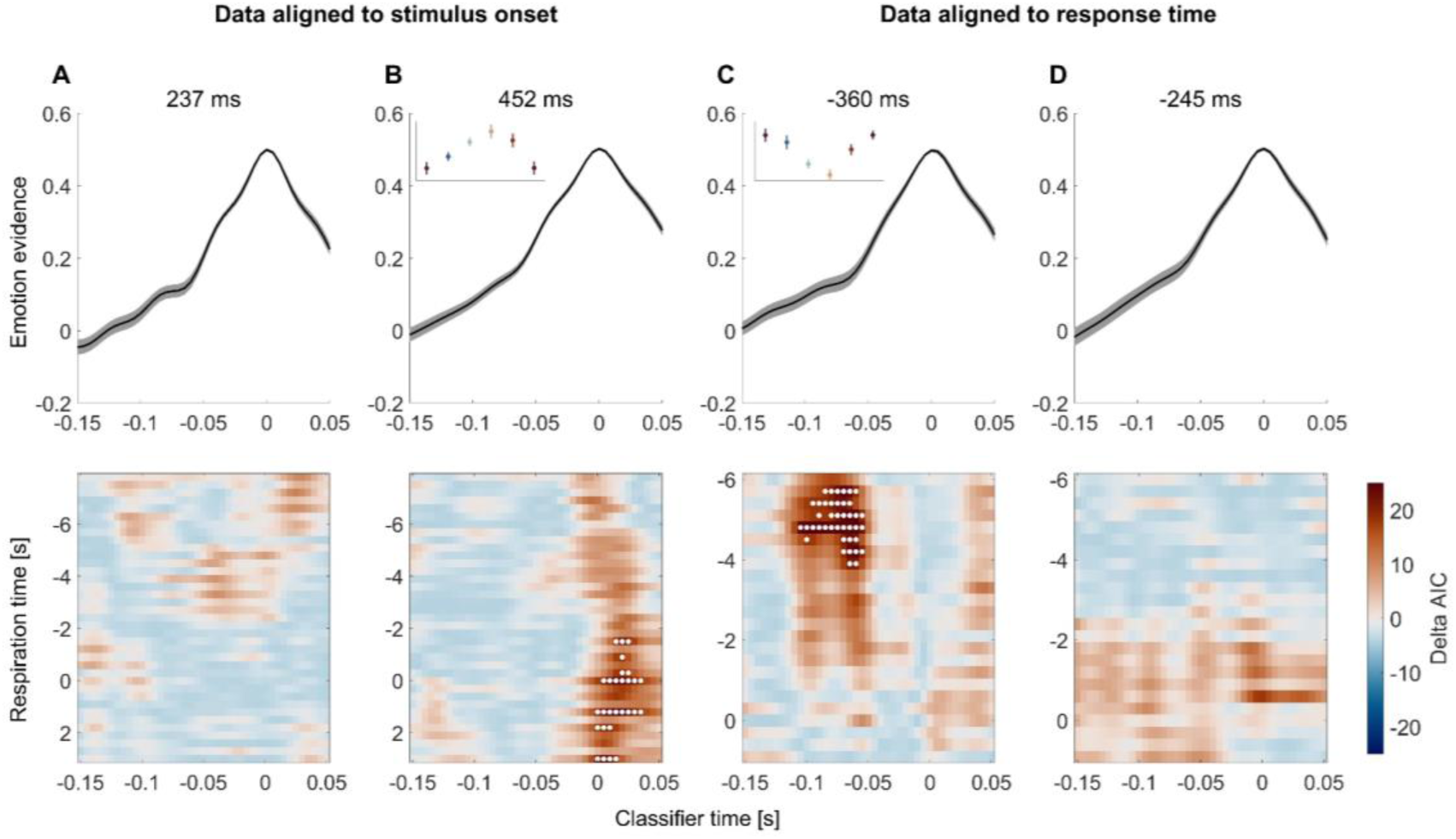
Modulation of classifier evidence by respiration. For each of the four EEG components (panel A-D) the upper panels show the emotion evidence carried by each component. Here classifier time refers to the time around the peak time at which the respective EEG component was defined in each participant. Black line shows group average with the standard error as grey shade. The lower panel shows the AIC difference for a modulation of this classifier evidenced by respiratory phase (with positive values indicating an improvement in model prediction when including respiration). Here respiration time refers to the time relative to stimulus onset (for panels A, B) or relative to the trial-wise response (panels C, D) at which the respiratory phase was derived. White dots indicate a difference in the deltaAIC corresponding to a probability of the model including respiratory phase to explain the data better at p<0.0001. Insets in panels B & C show the classifier evidence binned into six respiratory phase bins at the time of significant modulation (white dots in corresponding lower panels).

Figure 4 shows the ramping of emotion evidence for each component (upper panel) and the evidence for a modulation of this evidence by respiratory phase (as deltaAIC; lower panel). The late component derived from the stimulus aligned data was modulated by respiration around the peak time of this classifier (classifier time 0 s; corresponding to about 450ms from stimulus onset). This was evidenced by the respective AIC values (see Fig. 4, white dots in lower panel) and was confirmed by a cluster-based permutation test, which revealed a significant effect between -5ms and +35ms (corresponding to 447 ms to 487 ms from stimulus onset; at p<0.001). Similarly, the early component derived from the response aligned data was modulated by respiration. This is visible in the AIC values and a cluster-based permutation test revealed a significant effect between -110 ms and -50 ms prior to its peak, corresponding to about -460 to -400 ms prior to the response (cluster significant at p<0.001).

Having shown that cerebral signatures of the encoding of emotion category are modulated by respiration, we asked whether the sensory evidence carried by these components is also related to participants’ behaviour. For this, we probed an influence of the single-trial EEG evidence on behaviour on top of the influence of stimulus properties (emotion category, level) and respiration phase by themselves. For the late EEG component relative to stimulus onset this was indeed significant, with higher emotion evidence in the EEG leading to faster reaction times (beta = -0.006, p=0.00007, t=-3.99, deltaAIC=13.9) and higher response accuracy (beta=0.096, p=0.00004, t=4.12, deltaAIC=15.0). For the early EEG component in the response-aligned data this relation was not significant (RT: beta=0.001, p=0.64, t=0.474, deltaAIC=-1.8; accuracy: beta=0.014, p=0.53, t=0.627, deltaAIC=-1.6).

### These findings translate to the encoding of abstract visual stimuli

The above results show that respiration modulates the encoding of the sensory evidence about emotional faces. Using the data from the cube paradigm we show that this finding also translates to more abstract stimuli. For the shape paradigm, participants’ accuracy was influenced by the cube variant (p=0.006, t=2.74, deltaAIC = 5.5) and by the level of ambiguity (p<10^-5^, t=10.3, deltaAIC=112). The same was true for reaction times (cube variant: p<10^-5^, t=-12.65, deltaAIC=156; ambiguity: p<10^-5^, t=-10.25, deltaAIC=102; Table 1).

As for the emotion paradigm, participants’ respiratory phase was systematically structured around the individual trials (Fig 5). Again we observed a significant co-modulation of reaction times with respiration when aligning the data to the time of response (Fig. 5C, peak deltaAIC = 59.5, peak at -4 s) but not for response accuracy (Fig. 5D, peak deltaAIC = 9.33, peak at 0 s). When aligning the data to stimulus presentation the co-modulation was present for reaction times (peak deltaAIC = 18.5, peak at 1.8 s) but not present for accuracy (peak deltaAIC = 2.0, peak at -0.6 s).

**Figure 5:**
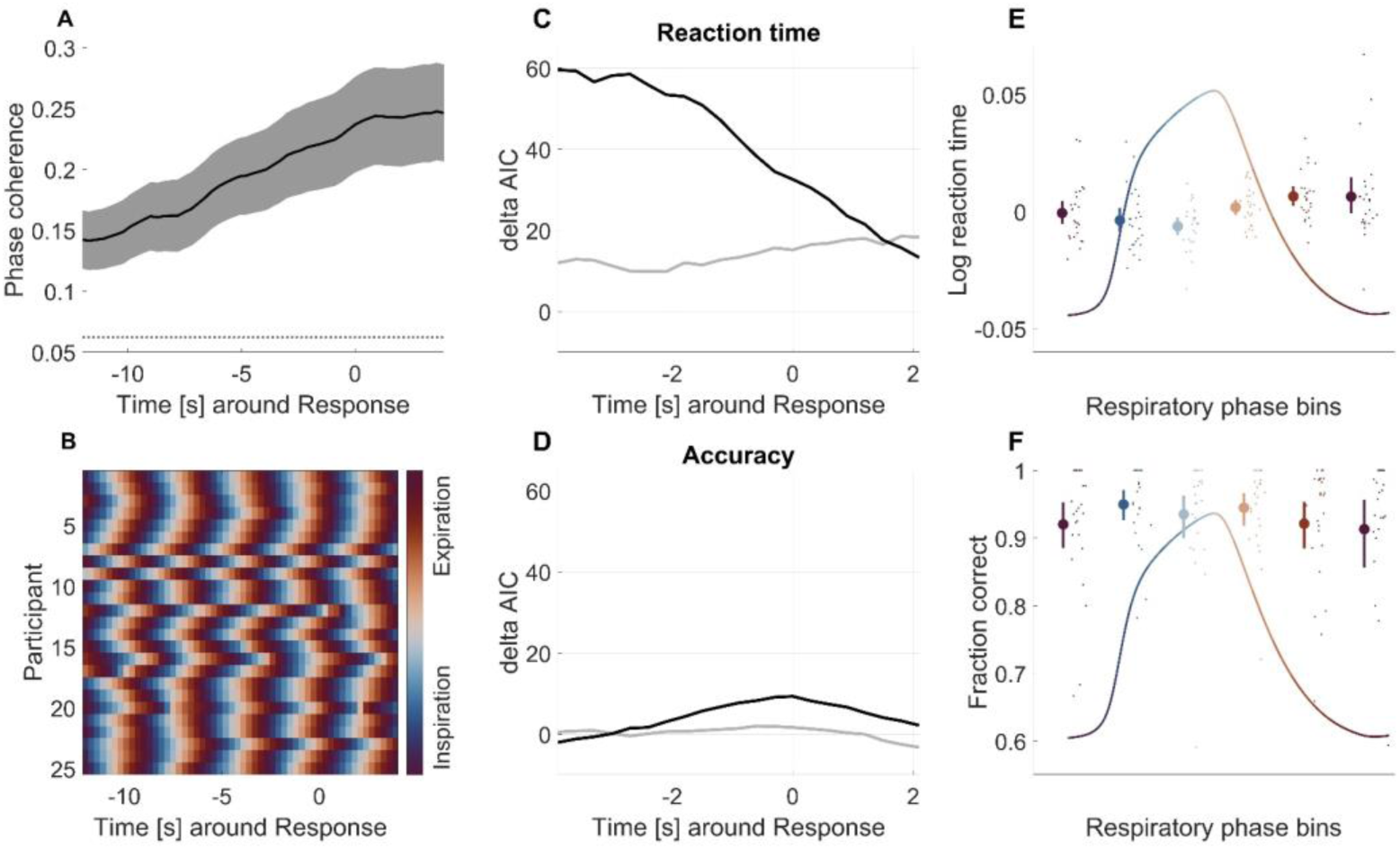
Behavioural and respiratory data for the shape paradigm. (A) Consistency of the respiratory phase across trials relative to the response times (mean, s.e.m); dotted line indicates significance threshold (p<0.01). (B) Trial-averaged respiratory phase for each participant. (C) AIC difference for a predictive effect of respiration on the reaction time; the grey line shows the effect for respiration aligned to stimulus onset, the black line for respiration relative to response times. Positive values indicate an effect of the respiratory phase. (D) Same for accuracy. (E) Average normalised reaction time (mean, ± s.e.m.) across participants for trials with a given respiration phase (divided into six bins; at the time of response). Dots indicate individual participants. (F) Accuracy across participants for each respiratory phase bin.

Using LDA we defined EEG components sensitive to the cube variant (Fig. 6). As for the emotion paradigm we derived two distinct EEG components during the time period of significant classification for both the stimulus- (peak times at 267 ms and 382 ms) and response-locked data (-440 ms and -255 ms). The topographies show a similar pattern as in the emotion paradigm with relative engagement of fronto-occipital electrodes. The average reaction time in this paradigm was around 680 ms which again suggests that the late component obtained relative to stimulus onset reflects similar processes as the early component obtained relative to response times.

**Figure 6:**
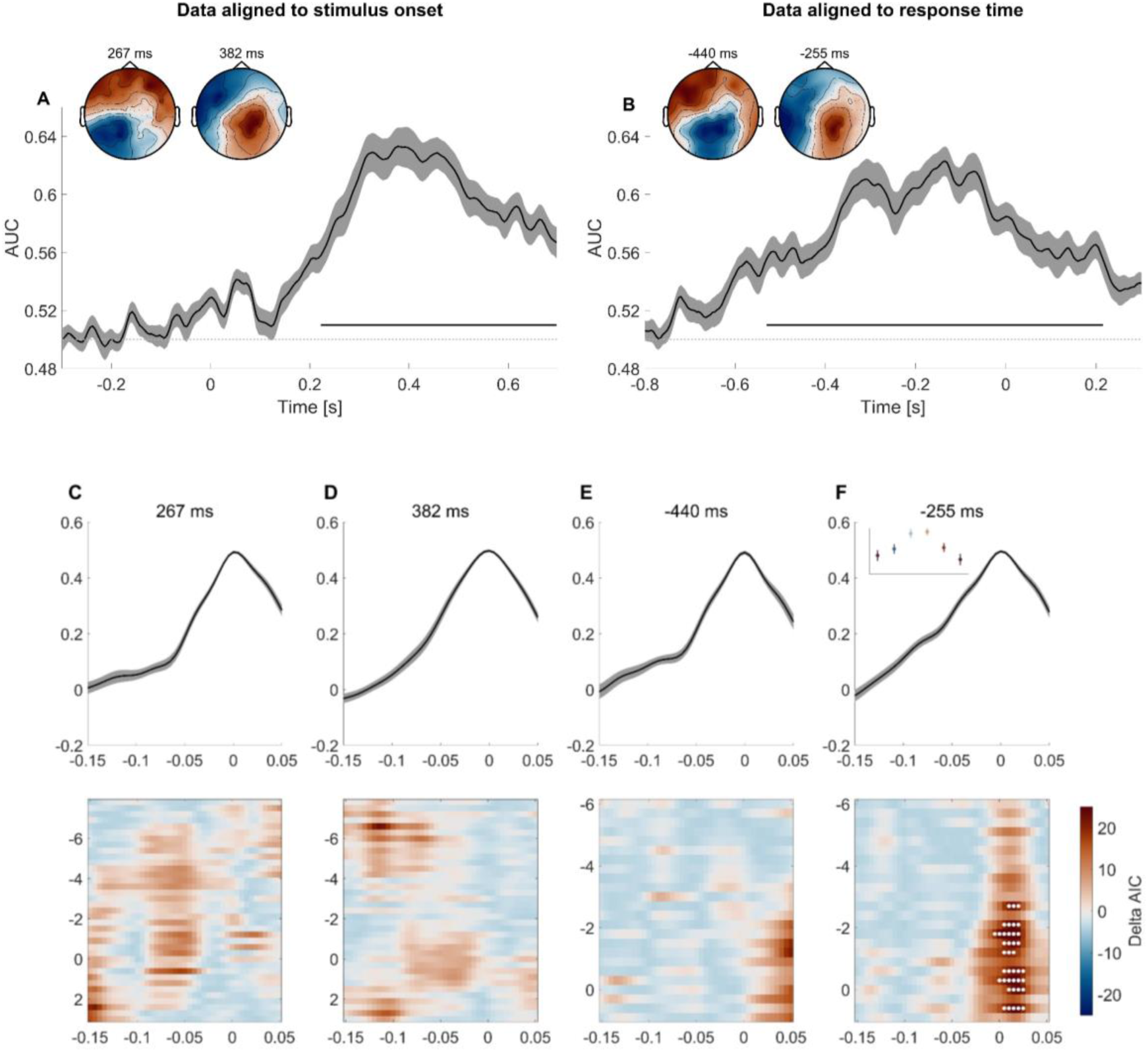
EEG components carrying evidence about the visual shape. (A) Classifier performance (AUC) for the data aligned to stimulus onset. Black line shows group average with the standard error as grey shade. Based on the LDA forward models we identified two components, the first peaking around 267 ms, the second around 382 ms after the stimulus onset. (B) Classifier performance for the data aligned to the response. Again two components were identified, one peaking 440 ms before the response, the other 255 ms before the response. The black horizontal lines indicate significant classifier performance (at p<0.01; cluster-based permutation test). (C-F) Modulation of classifier evidence by respiration. For each of the four EEG components the upper panels show the emotion evidence carried by each component. Here classifier time refers to the time around the peak time at which the respective component was defined in each participant. The lower row shows the evidence for a modulation of this evidence by the respiratory phase (AIC difference of the linear regression model with and without respiratory phase). Here respiration time refers to the relative to the stimulus onset (for panels C, D) or relative to the trial-wise response (panels E, F) at which the respiratory phase was derived. White dots indicate a difference in the deltaAIC corresponding to a probability of the model including respiratory phase to explain the data better at p<0.0001. Inset in panels D shows the classifier evidence binned into six respiratory phase bins at the time of significant modulation (white dots in corresponding lower panel).

As for the emotion paradigm we found a modulation of the classifier evidence by respiration (Fig. 6). Here this was present for the late component relative to the response and visible both in the AIC values and confirmed using a cluster-based permutation test. The latter showed a significant effect from -10 ms to 30 ms around the classifier peak (p<0.001). As for the emotion paradigm, the single trial sensory evidence of this respiration-modulated EEG component was predictive of participants’ behaviour, with stronger classifier evidence resulting in faster reaction times (beta=-0.006, p<10^-5^, t=-4.84, deltaAIC=21.4) and better response accuracy (beta=0.213, p<10^-5^, t=5.76, deltaAIC = 31.6).

## Discussion

The aim of this study was to investigate the neurophysiological basis of a co-modulation of behaviour with the respiratory cycle. Across two experiments we show that participants align their respiration to the experimental trials, while between-trial variations in respiratory phase are predictive of the behavioural outcome. Importantly, the respiratory phase modulates the encoding of task-relevant neurophysiological representations of the visual stimuli and those respiration-modulated cerebral signatures are predictive of behaviour. These results show that respiration not only has a modulatory influence on large-scale brain activity, but directly modulates task-specific neurophysiological processes underpinning behaviour.

### The respiratory phase modulates how neurophysiological signals reflect external sensory information

Many studies show that how quick or accurate participants respond in sensory or cognitive tasks depends on their respiratory phase during and prior to each trial (Flexman et al., 1974; Gallego et al., 1991; Huijbers et al., 2014; Arshamian et al., 2018; Perl et al., 2019; Johannknecht & Kayser, 2022). Yet, the precise neurophysiological mechanisms underlying this effect remain unclear. The present study corroborates the relation between respiration and behavioural performance and suggests that this arises from the specific influence of respiration on neurophysiological processes reflecting the encoding of task-relevant information used to guide perceptual judgements. This suggests a more specific influence of respiration on sensory-cognitive processes than known from previous studies.

Several studies have shown that signatures of rhythmic brain activity, such as delta, alpha or gamma oscillations are modulated by respiration across the brain. Studies in rodents, for example, show that respiration modulates oscillations in sensory cortices not involved in olfaction (Ito et al., 2014) as well as in parietal and prefrontal regions (Tort et al., 2018; Mofleh, et al., 2021; Jung et al., 2023) or shapes the functional connectivity of these with the limbic system (Mofleh et al., 2021; Liu et al., 2017). Such findings are also supported by direct intracranial recordings in human patients at rest, which show that the respiratory rhythm entrains local field potentials in the limbic system (Zelano et al., 2016) and oscillations in a widespread network of parietal and frontal areas (Herrero et al., 2018). Converging insights also come from human neuroimaging studies using MEG or EEG. These have shown that these neurophysiological signals vary in a coherent manner with respiration (Watanabe et al., 2023), resulting in a systematic co-modulation of the multiscale spectro-temporal organisation of brain activity with respiration in humans at rest (Kluger & Gross, 2021). Importantly, this includes markers of attention such as alpha band oscillations (Hsu et al., 2020; Kluger et al., 2021). Such a modulation of attention-related processes, or the neural excitation-inhibition balance (Kluger et al., 2023), may directly translate into a corresponding modulation of behaviour with the respiratory phase. However, the neurophysiological processes generating these large-scale signals are likely to be task-generic and do not imply that respiration necessarily modulates the quality by which sensory signals are encoded. In fact, a modulation of behaviour along the respiratory cycle could also originate from a direct influence of respiration on the motor system. In support of this, recent studies showed that the cortico-muscular coherence over the motor cortex (Kluger & Gross, 2020) and the motor readiness potential are shaped by respiration (Park et al., 2020). The present data extend this body of work by showing that respiration directly affects the quality by which brain activity represents task-relevant signals used to guide subsequent perceptual choices.

Related evidence has been obtained in studies on memory and learning. The quality of memory formation was found to vary with the phase of respiration during the encoding stage during image learning (Zelano et al., 2016; Johannknecht & Kayser, 2022), eye blink conditioning (Waselius et al., 2022) and a delayed-matched to sample task (Nakamura et al., 2018, 2022). At the level of brain activity these studies show that respiration modulates evoked responses to the conditioned stimuli and modulates fMRI-derived activity in parieto-frontal networks involved in memory (Waselius et al., 2022; Nakamura et al., 2022). The present data show that such a respiration-driven modulation is more widely present also during perceptual tasks not requiring learning and extends from generic evoked responses to task-related representations of external signals.

### The respiration-modulated EEG components presumably reflect decision processes

The timing and topographies of the respiration modulated EEG components give insights about the underlying processes that are modulated by the respiration phase. Previous studies have shown an influence of respiration on brain activity in early sensory areas (Waselius et al., 2022), in regions related to motor preparation and in brain-muscle coherence (Kluger & Gross, 2020) as well as in brain regions linked to decision processes (Biskamp et al., 2017; Zhong et al., 2017; Tort et al., 2021; Nakamura et al., 2022; Jung et al., 2023). The topographies of the relevant EEG components exhibiting an influence of respiration in the present study suggest a relative contribution of fronto-parietal electrodes in a bilateral manner. Furthermore, for both paradigms was the respiratory modulation stronger in EEG components emerging later in the trial and closer to the response than to stimulus onset: for the emotion paradigm the effect emerged between 353 to 313 ms prior to the response (based on the mean reaction time) and in the cube paradigm 265 ms to 224 ms prior to the response. Collectively, this points to activity in parietal or frontal regions that underlies the decision process as likely sources for the respective EEG components. While this does not rule out that information about the respiratory cycle is also present in early sensory regions, an origin in high-level brain regions is in line with the prominence of respiration-related signals in limbic, parietal and frontal brain regions. Studies using techniques with greater spatial resolution and focusing on signatures of sensory information coding as outlined here will be required to disentangle the respiratory influence on activity in early sensory and higher level brain regions.

### The function of body-brain interactions

Current theoretical frameworks on an influence of respiration on perception and cognition consider a central function in temporally structuring or organising large-scale neural processes (Heck et al., 2017; Brændholt et al., 2023; Krohn et al., 2023; Goheen et al., 2024; Kluger et al., 2024). Thereby respiration may contribute to shaping information encoding and communication in the brain. For example, MEG studies suggest that respiration modulates the excitation-inhibition balance and contributes to structuring rhythmic activity across regions (Kluger & Gross, 2023; Kluger et al., 2021, 2023). By contributing to temporally organising neural processes respiration may also facilitate the switch between an emphasis on external signals and an emphasis on interoceptive signals (Allen et al., 2023). In fact, the notion that bodily rhythms modulate sensory and cognitive processes may not only apply to respiration but also the heartbeat or even gut activity (Al et al., 2020, 2021, 2023; Engelen et al., 2024; Rebollo & Tallon-Baudry, 2022). For example, a study by Al and colleagues (2020) reported that during the systole of the cardiac cycle sensitivity for interoceptive signals is enhanced while during the diastole sensitivity is enhanced for external signals. Hence, various bodily rhythms may serve to predict or emphasise particular types of signals in order to facilitate the encoding and behavioural use of these.

